# Atypical processing of uncertainty in individuals at risk for psychosis

**DOI:** 10.1101/796300

**Authors:** David M. Cole, Andreea O. Diaconescu, Ulrich J. Pfeiffer, Kay H. Brodersen, Christoph D. Mathys, Dominika Julkowski, Stephan Ruhrmann, Leonhard Schilbach, Marc Tittgemeyer, Kai Vogeley, Klaas E. Stephan

## Abstract

**Background:** Current theories of psychosis highlight the role of abnormal learning signals, i.e., prediction errors (PEs) and uncertainty, in the formation of delusional beliefs. We employed computational analyses of behaviour and functional magnetic resonance imaging (fMRI) to examine whether such abnormalities are evident in at-risk mental state (ARMS) individuals.

**Methods:** Non-medicated ARMS individuals (*n*=13) and control participants (*n*=13) performed a probabilistic learning paradigm during fMRI data acquisition. We used a hierarchical Bayesian model to infer subject-specific computations from behaviour – with a focus on PEs and uncertainty (or its inverse, precision) at different levels, including environmental ‘volatility’ – and used these computational quantities for analyses of fMRI data.

**Results:** Computational modelling of ARMS individuals’ behaviour indicated volatility estimates converged to significantly higher levels than in controls. Model-based fMRI demonstrated increased activity in prefrontal and insular regions of ARMS individuals in response to precision-weighted low-level outcome PEs, while activations of prefrontal, orbitofrontal and anterior insula cortex by higher-level PEs (that serve to update volatility estimates) were reduced. Additionally, prefrontal cortical activity in response to outcome PEs in ARMS was negatively associated with clinical measures of global functioning.

**Conclusions:** Our results suggest a multi-faceted learning abnormality in ARMS individuals under conditions of environmental uncertainty, comprising higher levels of volatility estimates combined with reduced cortical activation, and abnormally high activations in prefrontal and insular areas by precision-weighted outcome PEs. This atypical representation of high- and low-level learning signals might reflect a predisposition to delusion formation.

## Introduction

In standard classification schemes such as the Diagnostic and Statistical Manual of Mental Disorders (DSM) and the International Statistical Classification of Diseases and Related Health Problems (ICD), schizophrenia and related psychotic disorders are defined as syndromes, i.e., combinations of clinical symptoms and signs. It is widely acknowledged that this phenomenological definition of schizophrenia (and other mental disorders) amalgamates heterogeneous groups of patients with possibly different pathophysiological mechanisms (van Os and Tamminga, 2007; Insel, 2010; Owen, 2014; Stephan et al., 2016). This may explain the diversity of clinical trajectories and treatment responses across patients and calls for new approaches to dissect the schizophrenia spectrum into subgroups or dimensions (Stephan et al., 2009a; Insel et al., 2010; Kapur et al., 2012; Cuthbert and Insel, 2013). One potential avenue to obtaining a formal understanding of differences in disease mechanisms across patients is the deployment of mathematical (specifically: generative) models that can be applied to non-invasive measures of behaviour and brain activity (Frässle et al., 2018); in psychiatry, the clinical application of this translational neuromodeling approach is referred to as “computational psychiatry” (Montague et al., 2012; Wang and Krystal, 2014; Stephan et al., 2015; Adams et al., 2016). A computational approach to phenotyping patients in a more fine-grained manner may be particularly relevant for the early detection of individuals at risk, an increasingly important domain of psychosis research (Klosterkotter et al., 2011; Fusar-Poli et al., 2013a, 2015; Moorhead et al., 2013; Koutsouleris et al., 2015; McGuire et al., 2015; Nieman and McGorry, 2015).

The at-risk mental state (ARMS) is defined by the presence of either attenuated psychotic symptoms, brief and self-limiting psychotic symptoms, or a significant reduction of function under a family history of schizophrenia (Fusar-Poli et al., 2013a). It is a construct pertaining to the pre-psychotic phase, before a formal diagnosis (according to DSM or ICD) can be made (Yung et al., 2005). Experimental studies and clinical trials of ARMS individuals are often logistically challenging, due to several reasons. For example, a diversity of strategies for diagnosis and clinical management exists across centres, and the unresolved question of whether ARMS symptoms mandate the use of antipsychotics (Wood et al., 2011; Kahn and Sommer, 2015; Nieman and McGorry, 2015) limits the availability of non-medicated ARMS individuals for research. Furthermore, not all ARMS individuals seek medical help and, if they do, they may not always be recognised due to frequent comorbidity, e.g., mood impairments may overshadow psychotic symptoms (Falkenberg et al., 2015a).

These issues lead to considerable difficulties in recruiting sufficiently large, homogeneous samples of medication-free (particularly antipsychotic-naïve) patients for clinical studies (Ruhrmann et al., 2010; Klosterkotter et al., 2011; Fusar-Poli et al., 2013a, 2013b; Nieman and McGorry, 2015). As a consequence, we still have a very limited understanding of the neurobiological mechanisms that produce ARMS symptoms and the subsequent transition to full-blown psychosis (Tsuang et al., 2013). A formal computational account of the cognitive and neurophysiological aberrancies underlying the ARMS would be highly beneficial, both for future research on pathophysiology and clinical studies.

In recent years, theories on the development and maintenance of psychotic symptoms have become increasingly enriched by testable computational mechanisms and are beginning to converge on a few central themes (Corlett et al., 2010). One major theory postulates that the attribution of “aberrant salience” to objectively uninformative or neutral events fuels the formation of delusions (Kapur, 2003). This framework posits a key role for dopamine (DA) in mediating the misattribution of salience in psychosis, consistent with longstanding theories of dopaminergic dysfunction in schizophrenia (Grace, 1993; Laruelle et al., 1996; Howes and Kapur, 2009) and specifically the idea that contextually inappropriate phasic DA release triggers maladaptive plasticity and learning (King et al., 1984; Heinz and Schlagenhauf, 2010; Roiser et al., 2013; Winton-Brown et al., 2014). This notion is consistent with a number of experimental findings in schizophrenia, most of which correlate with positive symptoms, including enhanced learning for neutral cues (assessed via behavioural and autonomic responses) and increased activation of dopaminergic and dopaminoceptive regions, including the ventral striatum and midbrain, in response to neutral cues (Jensen et al., 2008; Murray et al., 2008; Roiser et al., 2009; Romaniuk et al., 2010; Diaconescu et al., 2011).

Computational treatments of aberrant salience have examined this phenomenon in relation to prediction errors (PEs) about rewarding or novel outcomes. This is motivated by the putative relation of outcome-related PEs to phasic DA release and possible involvement in dysfunctional learning in psychosis and schizophrenia (Schultz et al., 1997; Pessiglione et al., 2006; Corlett et al., 2007, 2009a, 2010; Murray et al., 2008; Gradin et al., 2011; Adams et al., 2013; Ermakova et al., 2018). The dopaminergic PE signal is thought to represent a neural response to deviation from an expected outcome (of rewards but also sensory features; Iglesias et al., 2013; Gardner et al., 2018; Suarez et al., 2019) and likely supports updating of beliefs about the environment by induction of synaptic plasticity (Montague et al., 2004), for example, via modulation of NMDA receptors (Gu, 2002). However, predictions are inevitably uncertain, and PEs should carry different weight, depending on the precision (the mathematical inverse of uncertainty) of the underlying prediction. Computationally, Bayesian frameworks offer a formal account of this intuitive notion (Rao and Ballard, 1999; Friston, 2008). These theories view perception and learning as a hierarchically organised process, in which beliefs at multiple levels, from concrete (e.g., specific stimuli) to more abstract aspects of the environment (e.g., probabilities and volatility), are updated based on level-specific PEs and precisions. Specifically, under fairly general assumptions (i.e., for all probability distributions from the exponential family) a ratio of precisions (of bottom-up input vs. prior beliefs) serves to scale the amplitude of PE signals and thus their impact on belief updates (Mathys et al., 2011; Mathys, 2013); see Eq. 1 below.

Recent theories of perceptual abnormalities in schizophrenia have built on Bayesian accounts of this sort, enriching traditional concepts of aberrant salience with the crucial role of uncertainty (Stephan et al., 2006; Corlett et al., 2009a, 2010; Fletcher and Frith, 2009; Adams et al., 2013). One specific suggestion from these accounts is that chronically over-precise low-level PE signals may be the starting point of delusion formation, as they continue to induce unusual belief updates, without diminishing over time. Put differently, persistently surprising events may eventually require adoption of extraordinary higher-order beliefs to be explained away (Corlett et al., 2007, 2010). Alternatively, high-order beliefs may be of abnormally low precision (Adams et al., 2013; Sterzer et al., 2018; Diaconescu et al., 2019), leading to lack of regularisation, which renders the environment seemingly unpredictable (e.g., extremely volatile) and enhances the weight of low-level precision-weighted PE. Notably, these explanations are not exclusive but could co-exist (specifically, they relate to the numerator and denominator of the precision ratio in Eq. 1 below).

Here we investigated the presence of these putative abnormalities during learning and decision-making under environmental uncertainty (volatility) in the behaviour and brain activity of ARMS individuals. To this end, we combined fMRI of a probabilistic learning task under volatility with computational modelling, hierarchical Gaussian filtering (HGF), which emphasises the importance of uncertainty for updating a hierarchy of beliefs via precision-weighted PE signals (Mathys et al., 2011, 2014).

## Methods and Materials

### Participants

Thirteen ARMS individuals (mean age 21.1 years ± 3.0 s.d.; 4 female) and thirteen healthy controls (mean age 29.2 ± 3.2; 7 female) were included in the study. Given the slight differences in group composition, all statistical comparisons between the ARMS participants and controls in the fMRI analyses described below were conducted with age and sex as covariates, in order to control for possible confounding influences. Notably, none of the ARMS individuals had yet been exposed to any antipsychotic medications at the time of testing. Four of the ARMS individuals (2 female) had previously received medication for other mental health issues unrelated to psychosis. No ARMS individuals had any history of neurological disorder, and none of the healthy controls had any history of neurological or psychiatric disorder. A summary of demographic and clinical data is provided in Table 1. All participants provided written, informed consent to participate in the experiment, which was approved by the local ethics committee of the Medical Faculty of the University of Cologne (Cologne, Germany).

**Table 1.** Summary of demographic, behavioural and clinical variables (mean ± s.d.).

| | Measure | ARMS ($n = 13$) | Controls ($n = 13$) |
| --- | --- | --- | --- |
| Demographics | Age | 21.1 ± 3.0 | 29.2 ± 3.2 |
|  | Sex (M:F) | 9:4 | 6:7 |
| Behavioural parameters | $\kappa$ | 1.07 ± 0.20 | 1.13 ± 0.19 |
| | $\omega$ | -5.48 ± 1.74 | -5.05 ± 2.27 |
| | $\vartheta$ | 0.30 ± 0.04 | 0.31 ± 0.05 |
| | $\beta$ | 9.06 ± 4.26 | 7.03 ± 2.91 |
| | $m_3$ | 2.49 ± 0.36 | 2.21 ± 0.35 |
| | $\mu_3^{(k=0)}$ | 1.29 ± 0.57 | 1.80 ± 0.56 |
| Clinical variables | SIPS positive symptom sum | 6.85 ± 3.05 | |
| | SIPS negative symptom sum | 11.62 ± 3.66 | |
| | SIPS-GAF | 62.54 ± 10.14 | |
| | SIPS total | 31.23 ± 6.47 | |

The clinical status of ARMS individuals was established via a checklist of inclusion criteria and clinical tests including the Structured Interview for Prodromal Symptoms (SIPS; Miller et al., 2002, 2003). One male individual in the patient group did not have his ARMS status upheld upon further clinical examinations; his data were thus excluded from all between-group fMRI analyses. Scores on sub-scales of the SIPS assessing positive symptoms and global assessment of functioning (SIPS-GAF) were compared with data from the behavioural task and with a representative fMRI measure of aberrant PE encoding in ARMS, described below, via Pearson correlation analyses with one-tailed hypothesis testing, due to the explicit directionality of our predictions that atypical neural representation of PEs would be associated with worsened clinical functioning in ARMS.

### Behavioural task

While undergoing fMRI, each participant completed a probabilistic learning task (Fig. 1A), which required trial-wise binary decisions between two fractal stimuli. On each trial, the same pair of fractal stimuli were presented, each paired with a reward value (points) that varied independently of the task contingency structure. Accrued points were converted to monetary reward at the end of the experiment. During cue presentation, participants had a maximum of 4 seconds to make a decision, followed by a 5-second delay displaying the choice and a 2-second presentation of the decision outcome (correct or incorrect). If no response was made during the decision period, a time-out occurred and a blank screen was displayed for 8 seconds, instead of the delay and outcome screens. The inter-trial interval was jittered between 5 and 7 seconds. The across-trial task structure incorporated ‘switches’, or reversals, in a block-wise fashion in terms of which of the cues was most likely (80% probability) or least likely (20%) to be the correct, rewarded choice on that trial. It also incorporated a change of contingencies over time; this volatility (or variance per unit time) induces higher-order (environmental) uncertainty – about the probabilistic structure of the task, in addition to informational uncertainty by trial-wise outcomes – and determines the temporal evolution of a subject’s learning rate (Behrens et al., 2007; Mathys et al., 2014). The task consisted of 160 trials, with contingency blocks comprising between 14 and 46 trials depending on which volatility pseudo-block they were contained within (Fig. 1B). The task structure was fixed across all participants in terms of the correct, rewarded choice on a given trial. Two early participants (one patient and one control) completed a slightly varying version of the task that differed only in that there was no time-out for responses; this did not lead to longer data acquisition, nor did it affect the behavioural modelling described below.

**Figure 1.**
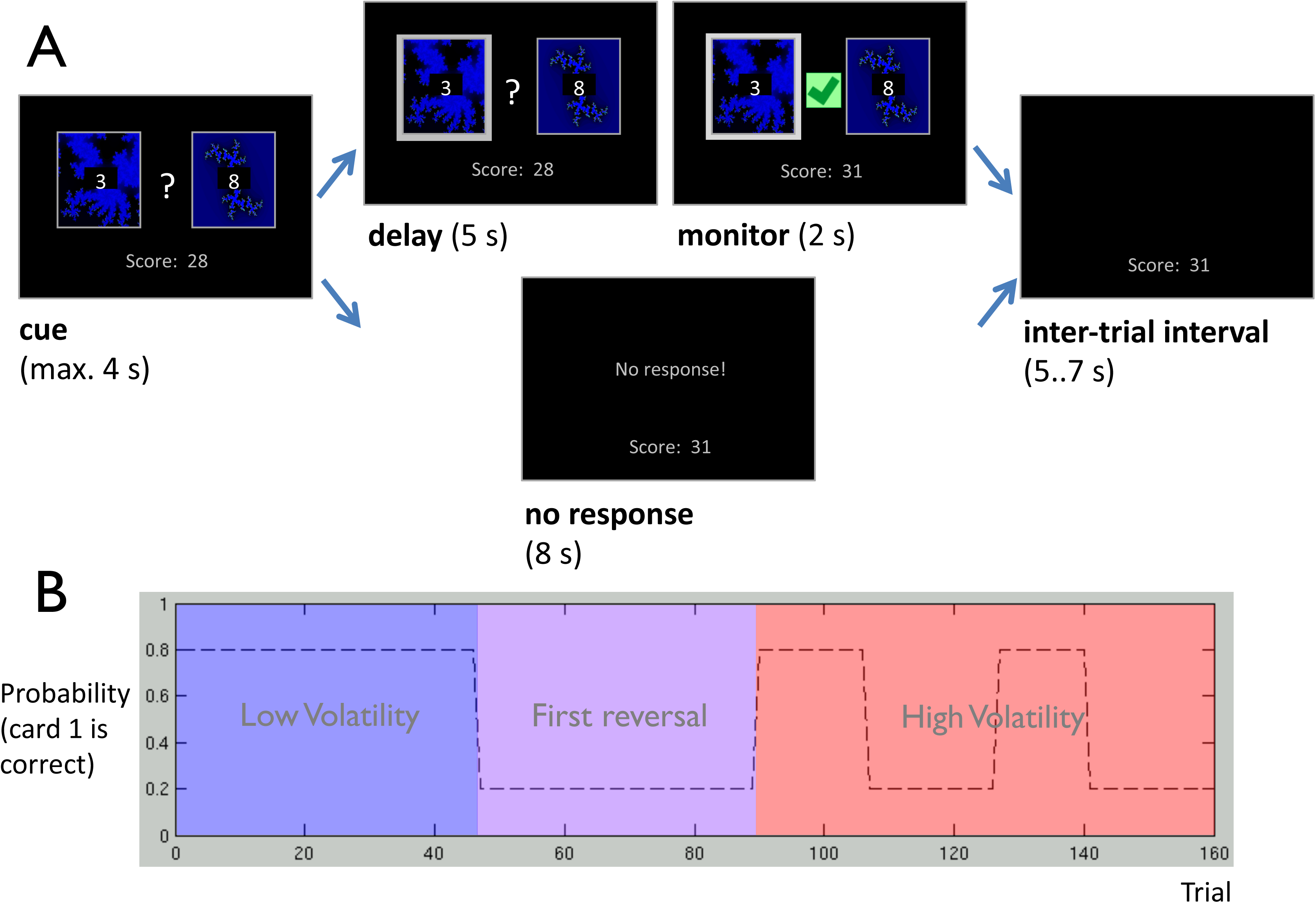
Probabilistic reversal-learning task. The behavioural paradigm consisted of: (A) within trial, a pair of fractal stimuli, each paired with a reward value, requiring a decision from the participant via button press in order to obtain the reward; (B) across trials, the probabilistic contingency (dotted line) of which of the two fractal cues was most likely to yield a reward occasionally underwent ‘reversal’, the regularity of which engendered pseudo-blocks of volatility modulation (blue, violet and red panels). The reward values within trials were entirely independent of the stimulus-outcome contingencies.

### Behavioural modelling

The computational framework adopted in this study was guided by Bayesian theories of brain function that suggest that the brain maintains and continuously updates a generative model of its sensory inputs (Dayan et al., 1995; Rao and Ballard, 1999; Friston, 2005). In other words, individuals are thought to update their beliefs about states of the external world based on the sensory inputs they receive (perceptual model); these beliefs, in turn, provide a foundation for making decisions (response model; see Daunizeau et al., 2010).

A number of different hypotheses about how humans learn about probabilistic stimulus-outcome contingencies were embodied in the following model space (Fig. 2A). With regard to the perceptual model, our main question was whether subjects’ models of reward probabilities based on stimulus-outcome associations had a hierarchical structure and accounted also for the volatility of these associations. We thus compared (i) the classical Rescorla-Wagner (RW) model (Rescorla and Wagner, 1972), in which predictions evolve as a function of PE and a constant learning rate (model *M*_1_) to three Bayesian models of learning, which included (ii) a non-hierarchical model, based on a reduced Hierarchical Gaussian Filter (HGF) that assumes that subjects do not infer on the volatility of stimulus-outcome probabilities (*M*_2_; see Diaconescu et al., 2014), (iii) a three-level ‘canonical’ HGF (*M*_3,4_; see Mathys et al., 2011), and (iv) a three-level ‘mean-reverting’ HGF in which volatility estimates drift towards a subject-specific equilibrium (*M*_5,6_).

**Figure 2.**
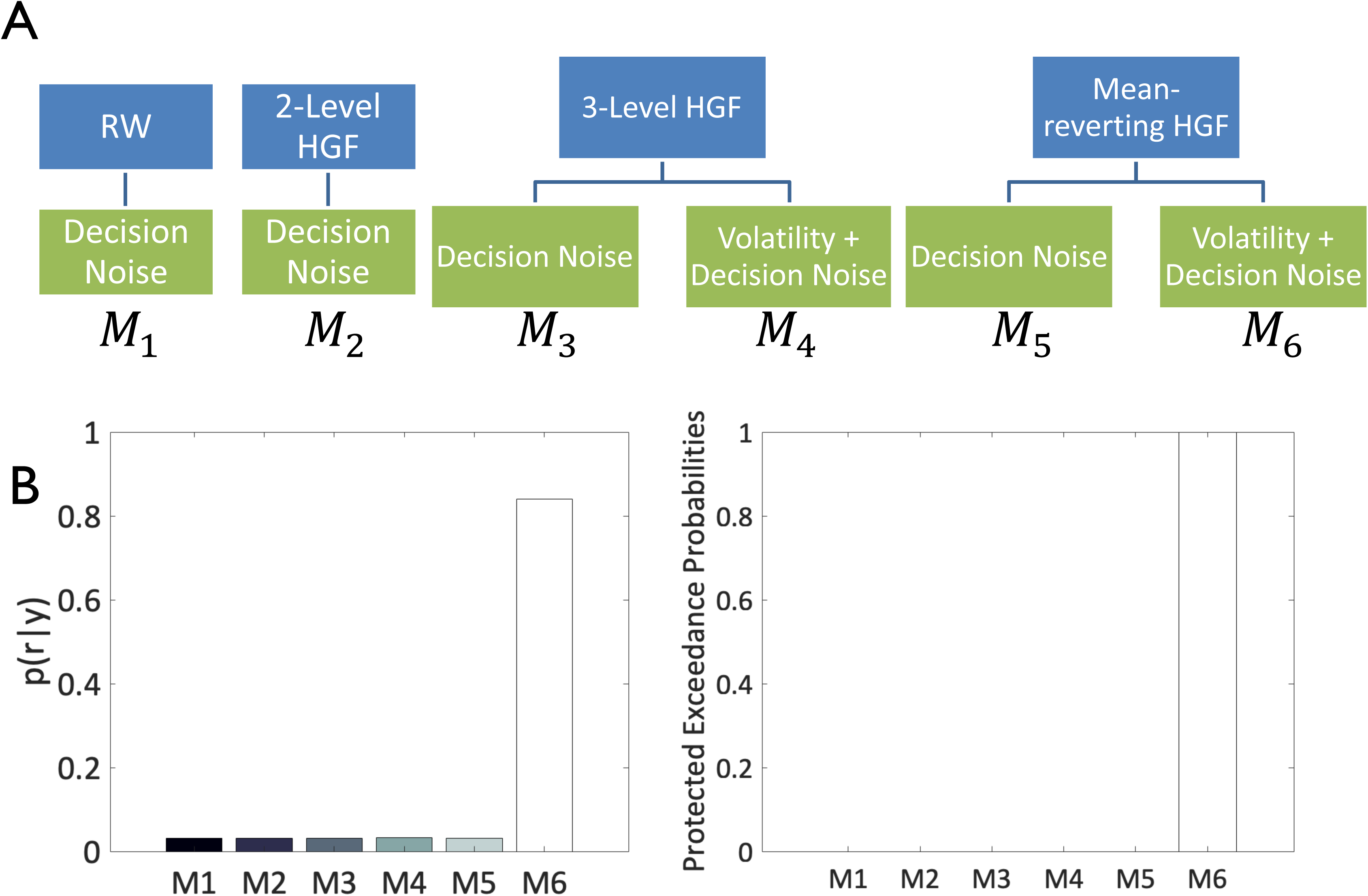
Hierarchical structure of the model space: perceptual models, response models and Bayesian model selection. (A) The models considered in this study have a factorial structure that can be displayed as a tree: The nodes at the first level represent the perceptual model families (RW, 2-level non-volatility HGF, 3-level HGF, and 3-level mean-reverting). The nodes at the second level represent the individual models. Two response model families were formalized under the HGF models: the mapping of beliefs-to-decisions either (i) depended dynamically on the estimated volatility of the learning environment (“Volatility + decision noise” model) or (ii) was a fixed entity over trials (“Decision noise” model). (B) Bayesian model selection (BMS) reveals *M_6_*, the mean-reverting HGF perceptual model in combination with the “Volatility” decision model, to best explain the data.

With respect to the response model, we followed previous work (Diaconescu et al., 2014) and considered two possible mechanisms of how beliefs were translated into responses (Fig. 2A). Subjects’ choices could either be affected by fixed decision noise (“Decision noise” model family; *M*_1-3,5_) or the decision noise might vary trial-by-trial as a function of the estimated volatility of the stimulus-outcome probabilities (“Volatility” model family; *M*_4,6_).

#### Perceptual model: the Hierarchical Gaussian Filter

The HGF is a hierarchical model of learning, which allows for inference on an agent’s beliefs (and their uncertainty) about the state of the world from observed behavior (Mathys et al., 2011) and has been used by several recent behavioural and neuroimaging studies on different forms of learning (Iglesias et al., 2013; Diaconescu et al., 2014, 2017; Hauser et al., 2014; Vossel et al., 2014; Schwartenbeck et al., 2015; Lawson et al., 2017; Powers et al., 2017). The model proposes that agents infer on the causes of the sensory inputs using hierarchically-coupled belief updates that evolve in time as Gaussian random walks where, at any given level, the variance (step size) is controlled by the state of the next higher level (Mathys et al., 2011, 2014). A standard formulation of the HGF for standard binary decision making tasks includes three levels, where the first (lowest) level encodes the probability of a trial outcome (here: whether a stimulus was rewarded or not), the 2^nd^ level represents the tendency of a stimulus to be rewarded as a continuous quantity (in logit space), and the 3^rd^ level represents the volatility of this probability (Fig. 3; see also Mathys et al., 2011).

**Figure 3.**
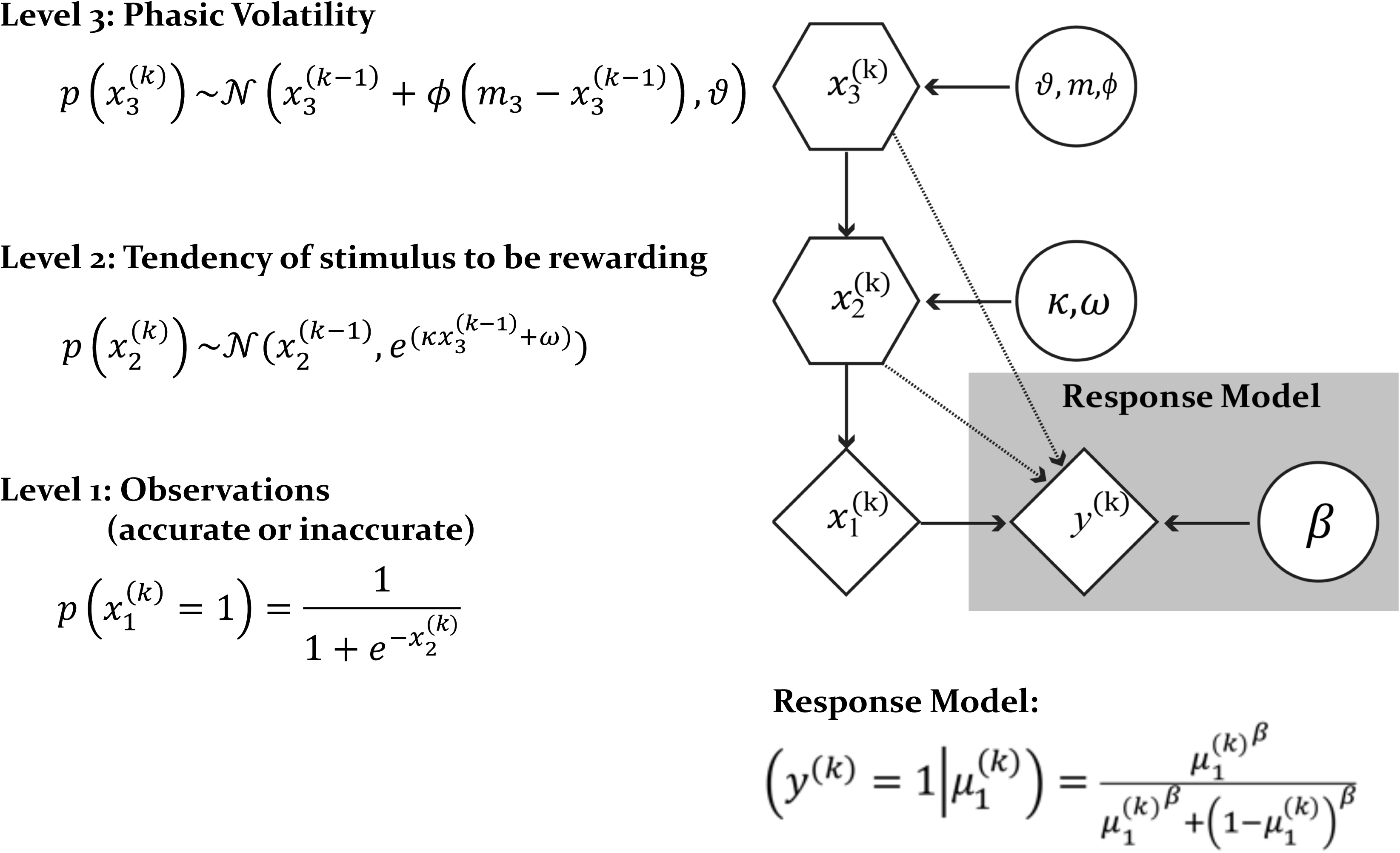
Graphical representation of the winning model combination: “mean-reverting HGF” perceptual model and the “Volatility” response model. In this graphical notation, circles represent constants and diamonds represent quantities that change in time (i.e., that carry a time/trial index). Hexagons, like diamonds, represent quantities that change in time, but additionally depend on the previous state in time in a Markovian fashion. x1 represents the cue probability, x2 the cue-outcome contingency and x3 the volatility of the cue-outcome contingency. Parameter κ determines how strongly x2 and x3 are coupled, ω determines the log-volatility or tonic component of x2, ϑ represents the volatility of x3, and m represents the mean of the drift towards which x3 regresses to in time. The response model parameter β represents the inverse decision temperature and determines the belief-to-response mapping.

The following subject-specific parameters determine how the above states evolve in time: (i) *κ* determines the degree of coupling between the second and third level in the hierarchy (*x*_2_ and *x*_3_) and the degree to which the volatility estimate influences the subject’s uncertainty about the stimulus-reward probabilities; (ii) *ω* represents a constant (tonic) component of the log-volatility of *x*_2_, capturing the subject-specific magnitude of the belief update about the stimulus-outcome probabilities that is independent of *x*_3_; (iii) *ϑ* is a meta-volatility parameter and determines the evolution of *x*_3_, or how rapidly the volatility of the associations changes in time. Furthermore, we also estimated 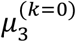, the subject’s initial belief about volatility of the outcome probabilities.

A key notion of the HGF is that subjects update their beliefs about hierarchically coupled states in the external world by using a variational approximation to intractable full Bayesian inference (Mathys et al., 2011). The update rules that emerge from this approximation have a structural form similar to RL, but with a dynamic (adaptive) learning rate determined by the next-higher level in the hierarchy. Formally, at each hierarchical level *i*, predictions (posterior means 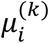) on each trial *k* are proportional to precision-weighted PEs, 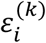 (Equations 1 and 2). The general form of this belief update (with subtle differences for categorical quantities at the lowest level) is the product of the PE from the level below 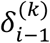, weighted by a precision ratio 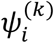:

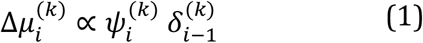

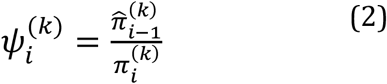

Here, 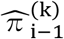 and 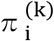 represent estimates of the precision of the prediction about input from the level below (e.g., precision of sensory data) and of the prediction at the current level, respectively (for details, see Mathys et al., 2011). This precision-weighting is critical for adaptive learning and emerges naturally from hierarchical Bayesian formulations (Friston, 2008; Corlett et al., 2010; Mathys et al., 2011; Adams et al., 2013; Iglesias et al., 2013). Simply speaking, PEs have a larger weight (and thus updates are more pronounced) when the precision of the data (input from the lower level) is high, relative to the precision of prior beliefs.

#### Mean-reverting HGF

The standard HGF, described above, already allows for representing (and inferring) the precision of low-level PEs and prior beliefs and thus offers the two components required to test our hypothesis. We can finesse this model further by using a variation of the classical HGF that allows for inferring on drifts in a subject’s beliefs towards an equilibrium *m* (essentially the equivalent of an Ornstein-Uhlenbeck process in discrete time). Here, we used this approach to examine the possibility that ARMS individuals might be characterised by a tendency to overestimate the volatility of the environment, which would further enhance the weight (precision) of low-level PEs and lead to an inflation of uncertainty about probabilities over time. As described above, a scenario of this sort may lead to later compensation, for example by adopting high-order beliefs with inappropriately high certainty, and may thus represent a risk factor for delusion formation.

The equations describing the generative model are summarised in Fig. 3. Notably, in this model, we infer on both the subject’s individual starting estimate of volatility, 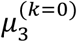, as well as a subject-specific parameter *m*_3_ (see Supplementary Table 1) that determines the equilibrium value to which the subject’s estimate of volatility drifts toward: the higher this value, the more uncertain the agent tends to be about his/her estimates of probabilities in the environment over time. The prior on *m*_3_ was equivalent to the prior on 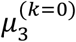, implying that over time, beliefs about volatility drift towards the agent’s prior. In other words, this model assumes that the agent has a tendency to ‘pull back’ estimates of volatility towards his/her prior expectations. The hypothesis described above, that ARMS individuals are characterised by enhanced precision-weighting of low-level PEs, implies that either the estimated precision of sensory input is abnormally high or that the precision of beliefs is abnormally low (see Eq. 2). In the context of our task, the latter corresponds to higher uncertainty about cue-outcome contingencies and implies that learning behaviour should be characterised by an upward drift of the predicted volatility, 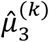, over time.

#### Response Model

The response model embodies a (probabilistic) mapping from the agent’s beliefs to their decisions (Daunizeau et al., 2010). The probability that the subject behaves according to his/her prediction of the outcome probabilities was described by a sigmoid function (Eq. 3):

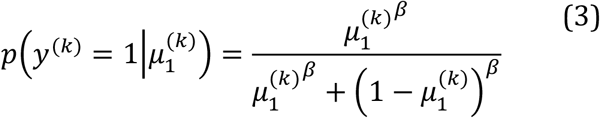

Here, *β* represents the inverse of the decision temperature: as *β* → ∞, the sigmoid function approaches a step function with a unit step at b^(k)^ = 0.5 (i.e., no decision noise). As described above, we considered two types of this belief-to-response mapping. The first “Decision noise” model family assumes constant decision noise; that is, *β* becomes a subject-specific free parameter. By contrast, the “Volatility” response model family proposes that the decision temperature parameter *β* might vary trial-by-trial with the estimated volatility, 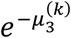, such that the larger the volatility, the lower the (inverse) decision temperature and the higher the decision noise (see Diaconescu et al., 2014 for details). In other words, the more stable the stimulus probabilities, the more deterministic the participant’s belief-to-response mapping.

Using priors over the model parameters based on previous studies with similar probabilistic learning paradigms (Iglesias et al., 2013; Diaconescu et al., 2014; Supplementary Table 1), maximum *a posteriori* (MAP) estimates of model parameters were obtained using the HGF toolbox version 3.0. This MATLAB-based toolbox is freely available as part of the open source software package TAPAS (https://www.tnu.ethz.ch/de/software/tapas.html).

#### Bayesian model selection and computational regressors

We compared the full set of resulting models *M*_1-6_ using Bayesian model selection (Stephan et al., 2009b), to determine which combination of perceptual and response models best explained the behavioural dataset and would thus optimally inform the subsequent analysis of fMRI data. Based on the model space outlined above (Fig. 2A), a total of six different models were compared.

From the winning model (Fig. 2B), we extracted the trajectories of several trial-wise computational quantities, estimated for each subject individually: (i) the prediction about the next outcome, (ii) uncertainty (Bernoulli variance) about the probability of the next outcome (‘1^st^-level uncertainty’), (iii) their belief about the current volatility of the environment 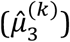, (iv) their absolute precision-weighted PE regarding the outcome on a given trial relative to their current beliefs about the probability of that outcome (*ε*_2_), (v) their belief uncertainty (‘2^nd^-level uncertainty’; *σ*_2_) and (vi) their signed precision-weighted PE regarding the perceived volatility of the outcome on a given trial relative to their current belief about that volatility (*ε*_3_). Each of these trajectories was then used as a regressor (parametric modulator) in the single-subject fMRI analyses described below.

### Behavioural analysis

We subjected the MAP estimates of the winning model to one-way analysis of variance (ANOVA) assessments, in order to test for differences in decision and learning parameters between ARMS individuals and healthy controls. Our hypothesis described above implies that *m*_3_ itself and/or the meta-volatility volatility parameter *ϑ* would be significantly greater in the ARMS group, suggesting that ARMS participants in contrast to controls perceive an increased environmental volatility over time. To examine group differences in perceived volatility induced by basic reversals of probabilistic contingency, we also performed a 2 (group: ARMS, control) × 3 (phase: stable, reversal, volatile) mixed-factor ANOVA to examine group-by-phase interaction effects on 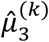 (see the phases outlined in Fig. 1B). The code for behavioural modelling and analysis is accessible via GitLab at https://gitlab.ethz.ch/dandreea/apup.

### Image acquisition

Data were acquired on a 3 T Magnetom TIM Trio MRI scanner (Siemens, Erlangen, Germany) at the Max Planck Institute for Metabolism Research, Cologne. As the task was partially self-paced, a slightly different number of volumes was acquired for each subject (mean = 1217, approximately 40.5 minutes experiment duration). T2*-weighted echo-planar images (EPI) sensitive to blood-oxygenation level-dependent (BOLD) contrast were acquired during the task (TR = 2 s; TE = 30 ms; flip angle = 90°; 30 axial slices; in-plane resolution = 3.3 × 3.3 mm; slice thickness = 2.7 mm; slice gap = 1.35 mm). Magnetic equilibration was accounted for via scanner automatic dummy removal. Images were acquired in parallel to the anterior-posterior commissural plane. Cardiac and breathing rates were recorded peripherally during scanning. Anatomical T1-weighted volumes were also acquired for each subject (TR = 1.9 s; TE = 3.51 ms; flip angle = 9°; field-of-view 256 × 256 × 128; voxel size 1.0 × 1.0 × 1.0 mm).

### fMRI data preprocessing

Preprocessing of fMRI data was performed using FSL 5.0 (http://fsl.fmrib.ox.ac.uk/fsl/fslwiki/; Centre for Functional Magnetic Resonance Imaging of the Brain, University of Oxford, United Kingdom; Smith et al., 2004). This incorporated standard steps of high-pass filtering (128 seconds cut-off), realignment of individual volumes to correct for head motion, removal of non-brain tissue from the images and Gaussian smoothing at 5 mm full-width half-maximum.

Following these initial steps, we performed single-subject data ‘denoising’: artefact classification and rejection based on single-session spatial independent component analysis (ICA) and machine learning techniques, using the FSL ‘MELODIC’ and ‘FIX’ tools (Beckmann and Smith, 2004; Salimi-Khorshidi et al., 2014). These steps were incorporated to remove any artefactual signal components that had survived conventional realignment and physiological noise correction (where available; see next section). Details are provided, along with a description of spatial normalisation procedures, in the Supplementary Materials.

### fMRI analysis

Single-subject fMRI analyses were conducted using the general linear model (GLM) as implemented in SPM8 (http://www.fil.ion.ucl.ac.uk/spm/software/spm8/; Wellcome Trust Centre for Neuroimaging, University College London, United Kingdom). Base regressors for the task were defined in terms of the onsets of the decision period, which had a variable duration (0-4 s) and the outcome period, which had a fixed duration (2 s). The decision period regressor was accompanied by three parametric modulator regressors encoding for the subject’s trial-wise prediction of outcome, uncertainty at the 1^st^ level of the HGF, and expected volatility at the 3^rd^ level. The outcome period regressor was associated with three parametric modulators encoding for the absolute (unsigned) outcome-related precision-weighted PE (ε_2_; (see Iglesias et al., 2013), uncertainty at the 2^nd^ level (*σ*_2_) and the volatility-related precision-weighted PE (ε_3_). All parametric modulators were Z-normalised (zero mean, unit standard deviation) before entering into the GLM. Orthogonalisation was not performed. Temporal and dispersion derivatives of all regressors were added to the GLM of each subject in order to account for variability in the onset and width of hemodynamic responses. We performed physiological noise correction, based on RETROICOR (Glover et al., 2000) of cardiac and respiratory measurements (where available; 8 controls and 7 patients), using the PhysIO toolbox in TAPAS (Kasper et al., 2017) to compute 18 additional regressors that were included, along with six realignment parameters representing head motion, as confound regressors in the GLM for each subject.

Group analyses were conducted using second-level GLMs as implemented in SPM8. Outcome-related contrast estimates from the subject-level analysis were entered into a 2 (diagnostic groups: ARMS vs. controls) × 4 (outcome-related variables: base regressor, *ε*_2_, *σ*_2_ (2^nd^-level uncertainty), *ε*_3_) analysis of covariance (ANCOVA; unequal variance assumed). Age and sex were included as covariates of no interest. Decision-related contrast estimates were entered into a similar 2 (group: ARMS vs. controls) × 4 (decision-related variables: base regressor, prediction, 1^st^-level uncertainty, volatility estimate) analysis of covariance (ANCOVA).

Contrasts of interest at the group level examined, for each computational regressor, (i) the average activation across groups (ARMS + controls) and (ii) significant differences between groups (ARMS vs. controls). For the latter analyses, we conducted whole-brain comparisons but also used contrast-masking to restrict the analysis to regions showing significant average effects across groups (note that these are orthogonal contrasts, thus avoiding problems of non-independent inference). One exception to the latter approach was made in the analysis of group differences in *ε*_3_, where (iii) we restricted analyses to an anatomically defined mask of the anterior portion of the cingulate cortex (defined using the probabilistic volume in the Harvard-Oxford atlas provided with FSL and further masked by a study-specific group-level EPI template). This *a priori* mask, which included the basal forebrain, was motivated by previous observations of *ε*_3_-related activation in the basal forebrain and anterior cingulate cortex (Iglesias et al., 2013; Diaconescu et al., 2017; see also Behrens et al., 2007). In line with the results of the same studies, we also employed (iv) a similar *a priori* region of interest analysis of *ε*_2_- and *ε*_3_-related activations using a mask of the dopaminergic midbrain. For each contrast, we corrected for multiple comparisons across the respective search volume – i.e., whole brain for (i), functional masks for (ii), and the anatomical mask for (iii) and (iv) – using family-wise error (FWE) correction at the cluster-level (p < 0.05, with a cluster-defining threshold of p<0.001).

### fMRI association with clinical variables

We hypothesised that key clinical features of the ARMS might be associated with the representation of low-level precision-weighted PEs in brain regions that were (i) activated by these learning signals and (ii) exhibiting aberrantly greater such activation in ARMS relative to healthy control participants. To this end, and in line with previous convergent findings indicating a link between prefrontal cortical brain changes and ARMS (Benetti et al., 2009; Allen et al., 2012; Cannon et al., 2015), we selected a prefrontal region comprising a cluster of differential fMRI activation to ε_2_ (see Results) in which to examine fMRI contrast beta-value associations (via Pearson correlation) with SIPS positive symptoms and SIPS-GAF measures, predicting that higher ε_2_-related activation in this region would be associated with greater symptom burden/severity.

## Results

### Behaviour

Initially, we applied a mixed-factor ANOVA with between-subject and within-subject factors to directly measurable behaviour (accuracy and reaction times, respectively) in order to test for significant main effects or interactions (2×3 factorial design: group × task phase). No main effects of group and no group × phase interactions were found. For performance accuracy, we observed a main effect of phase (df = (2, 48), F = 5.59, p = 0.008 with Greenhouse-Geisser (GG) nonsphericity correction, effect size 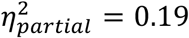), with reduced performance in volatile compared to stable phases of the task. By contrast, the main effect of phase for RT closely failed to reach significance (df = (2, 48), F = 3.11, p = 0.07 with GG correction, 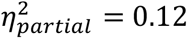). We then proceeded to computational analyses to test for group differences in latent variables underlying learning and decision-making.

### Behavioural modelling

Bayesian model selection gave a clear result, showing that the mean-reverting HGF with a response model incorporating volatility mapping (*M*_6_) was more likely to explain task behaviour than any other model type (Fig. 2B). A summary of parameters from the inversion of this winning model is provided for both groups in Table 1. Importantly, model selection results were equivalent in both groups, allowing for a direct comparison of parameter estimates across groups. Group average belief-updating trajectories computed from the winning model *M*_6_, are depicted in Figs. S1 and S2.

We investigated group differences in the parameters that (i) were well recovered from simulations and (ii) particularly impacted learning about volatility and thus the precision of high-order beliefs. While we did not observe any group differences for the meta-volatility *ϑ* parameter (df = (1, 25), F = 0.54, p = 0.47, 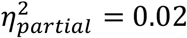), we found a significant difference in the mean-reversion equilibria values *m*_3_. Relative to controls, ARMS individuals exhibited reversion to significantly higher equilibria levels (group: df = (1, 25), F = 4.29, p = 0.049, 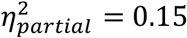; Fig. 4A). This effect remained significant, whether or not the single participant in the ARMS group whose diagnosis failed to be upheld at follow-up (and who was thus excluded from the between-group fMRI analyses) was included. From Figs. S1 and S2, one can observe that, in contrast to controls, the belief trajectories for the estimated volatility extend to higher levels in ARMS compared to controls, consistent with the group differences in the MAP estimates of *m*_3_.

**Figure 4.**
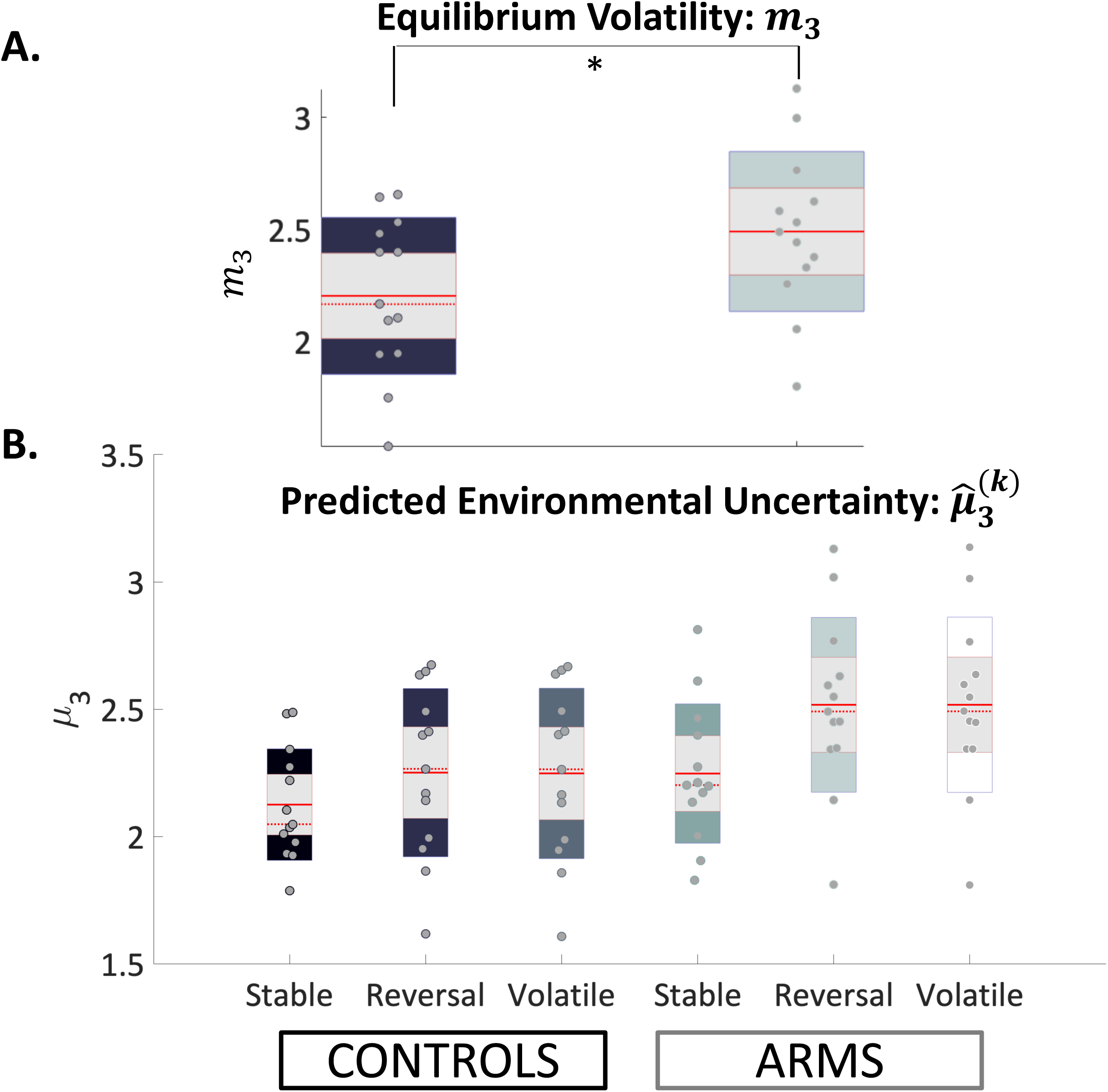
Behavioural parameter group differences. (A) Group differences in reversion equilibria values (*κ*_3_): larger reversion equilibria values were detected in ARMS compared to controls (group: df = (1, 25), F = 4.29, p = 0.049, 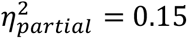); and (B) Group-by-phase interactions of perceived environmental volatility (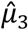): A mixed-factor ANOVA (with Greenhouse-Geisser nonsphericity correction), which included between-subject and within-subject factors, found a significant main effect of phase and a significant group = phase interaction (phase: df = (2, 48), F = 41.68, p = 1.113e-06, 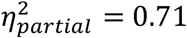; group x phase: df (2, 48), F = 5.71, p = 0.025, 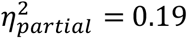). See main text for details. Jittered raw data are plotted for each parameter. The solid red line refers to the mean, the dotted red line to the median, the grey background reflects 1 SD of the mean, and the coloured bars the 95% confidence intervals of the mean. ‘*’ refers to group differences of significance level p < 0.05.

Second, we investigated group × task phase interaction effects on the phase-specific averages of predicted environmental volatility, or 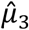, following probabilistic contingency reversals, by performing a mixed-factor ANOVA with between-subject and within-subject factors (with GG nonsphericity correction). We found a significant group-by-phase interaction effect (df = (2, 48), F = 5.71, p = 0.025, 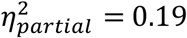; Fig. 4B), suggesting that, in comparison to controls, ARMS individuals display a significantly larger increase in perceived environmental volatility following the first probability reversal. We also found a main effect of phase (df = (2, 48), F = 41.68, p < 0.001, 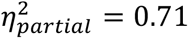), reflecting significant increases in perceived volatility for both groups with increases in the volatility of the task schedule. Importantly, these mixed-factor ANOVA results remained significant, whether or not the single participant in the ARMS group whose diagnosis failed to be upheld at follow-up was included.

A number of our model parameters were estimated based on participants’ choice behaviour. We also examined whether their actual values could be recovered from synthetic data. Thus, we simulated responses based on all participants’ MAP parameter estimates, and then fitted the model to those simulated responses. The results of the parameter recovery are included in Fig. S3. Several parameters could be recovered well from the data: The second-level parameters encoding tonic (*ω*) and phasic (*κ*) aspects of volatility and third-level parameters determining the dynamics of the volatility trajectory, including the equilibrium point *m*_3_ and the meta-volatility parameter *ϑ* were recovered from the data well. By contrast, the initial value on the volatility 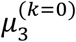, the initial variance on the volatility 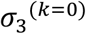, and the inverse decision noise parameter *β* could not be recovered from the data well.

### fMRI

When pooling across both groups, we found that the computational regressor associated with the precision-weighted outcome PE signal (*ε*_2_) activated a set of bilateral cortical regions (Fig. 5A), similar to previous analyses of the same type of PE from a sensory learning task (Iglesias et al., 2013). More specifically, the nine clusters forming this network were located bilaterally in inferior parietal cortex, anterior insula, ventrolateral prefrontal cortex (vlPFC) extending also into right dorsolateral prefrontal cortex (dlPFC) and superior frontal cortex (spanning superior and middle frontal gyri), as well as in a region of left cerebellum (p < 0.05, whole-brain cluster-level FWE-corrected, see Methods; Table 2). An additional region of interest analysis, using an anatomically defined *a priori* mask of the midbrain (Bunzeck and Düzel, 2006; Iglesias et al., 2013) revealed no significant activation in this region.

**Figure 5.**
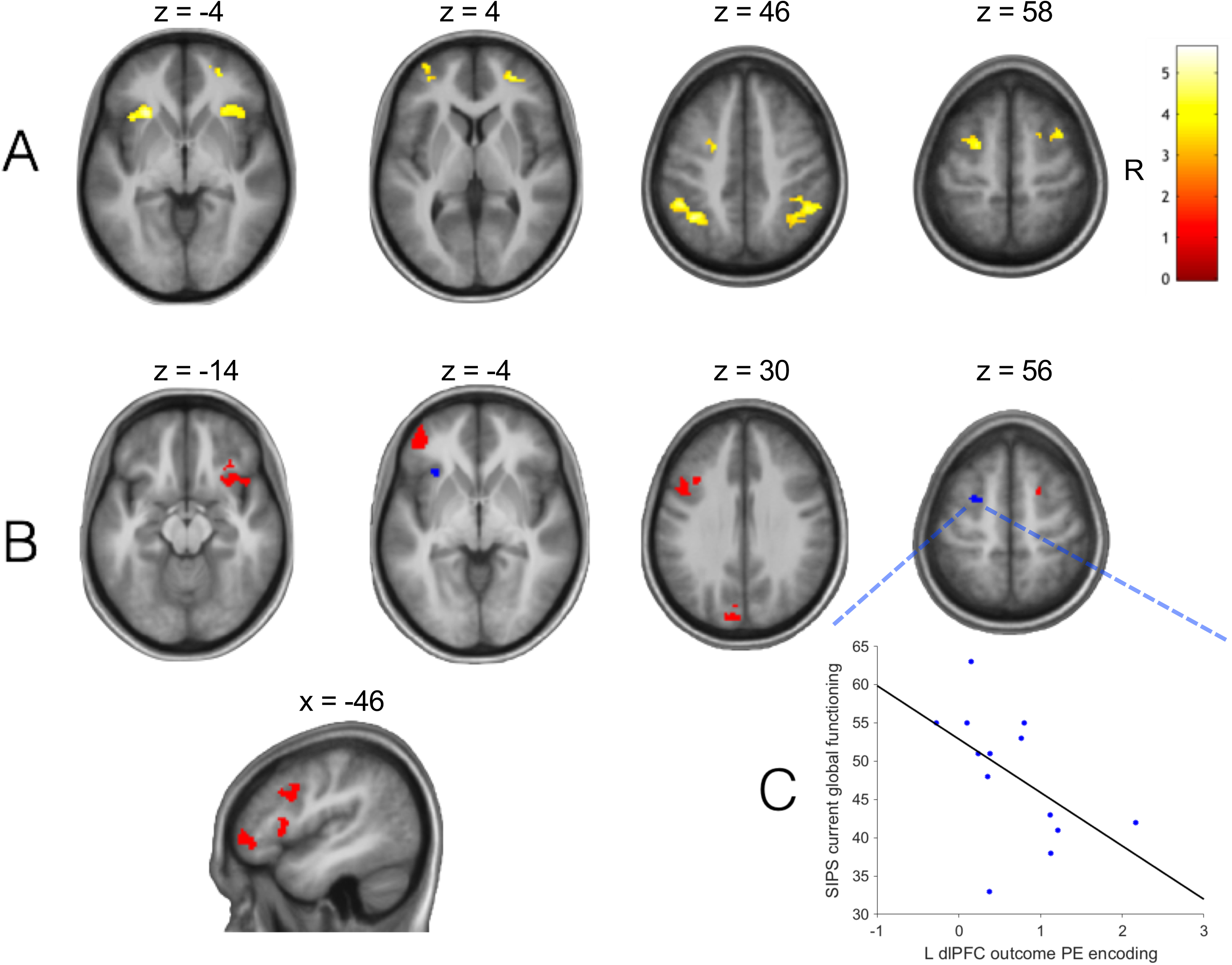
The neural representation of low-level/outcome-related precision-weighted PEs (*ε*_2_) in ARMS patients and healthy controls. (A) A representative map of significant (cluster-level FWE-corrected p < 0.05) group-level (ARMS + controls) outcome-related activations modulated parametrically by *ε*_2_, calculated via one-sample t-test (N = 25) and overlaid on an anatomical image calculated as the mean structural MRI of the whole group. (B) Significantly greater representation of *ε*_2_-related activation in a sub-set of these and other regions in ARMS patients relative to controls. Solid colour maps of group differences are binarised and indicate spatial differences between whole-brain cluster-level corrected results (red) and results corrected using the group average map in ‘(A)’ for contrast-masking (dark blue). Colour bar represents t-statistics. Axial and coronal slices are orientated in line with neurological conventions (R = right). (C) Significant negative correlation in ARMS patients (N = 13) between a clinical measure of current global functioning (SIPS-GAF) and *ε*_2_-related activation (beta-values, from the analysis as in panel ‘A’) in a region of left (L) superior dlPFC also showing significant group differences (ARMS > controls, from the independent analysis as in panel ‘B’).

**Table 2.**
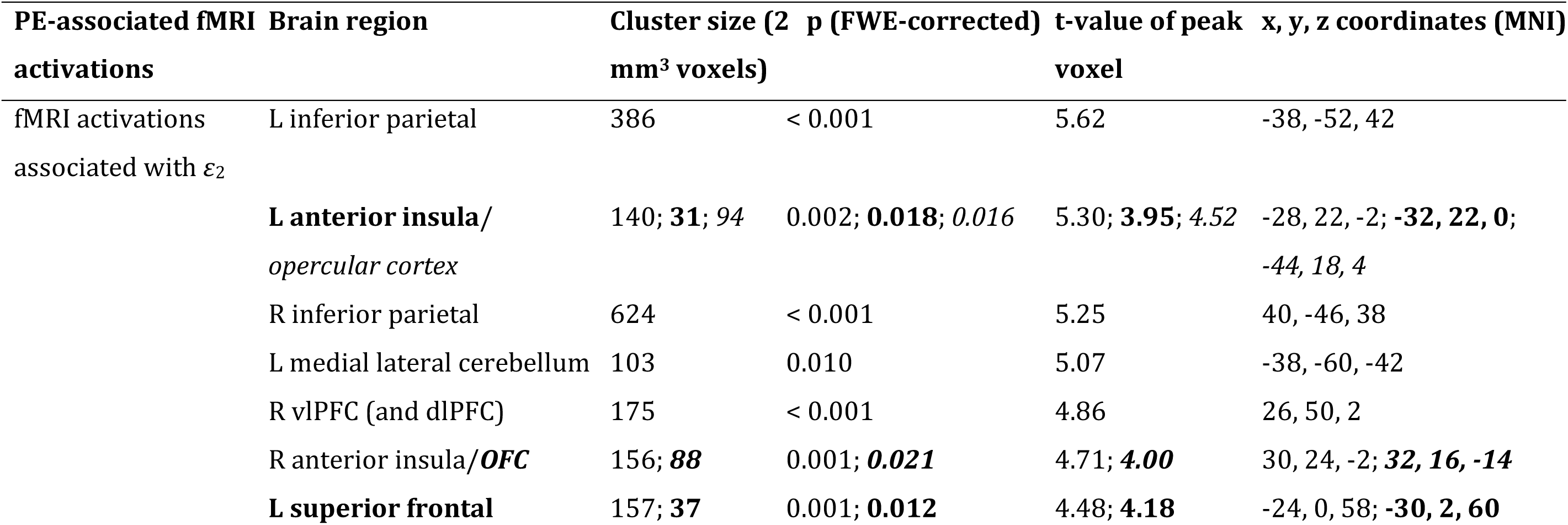
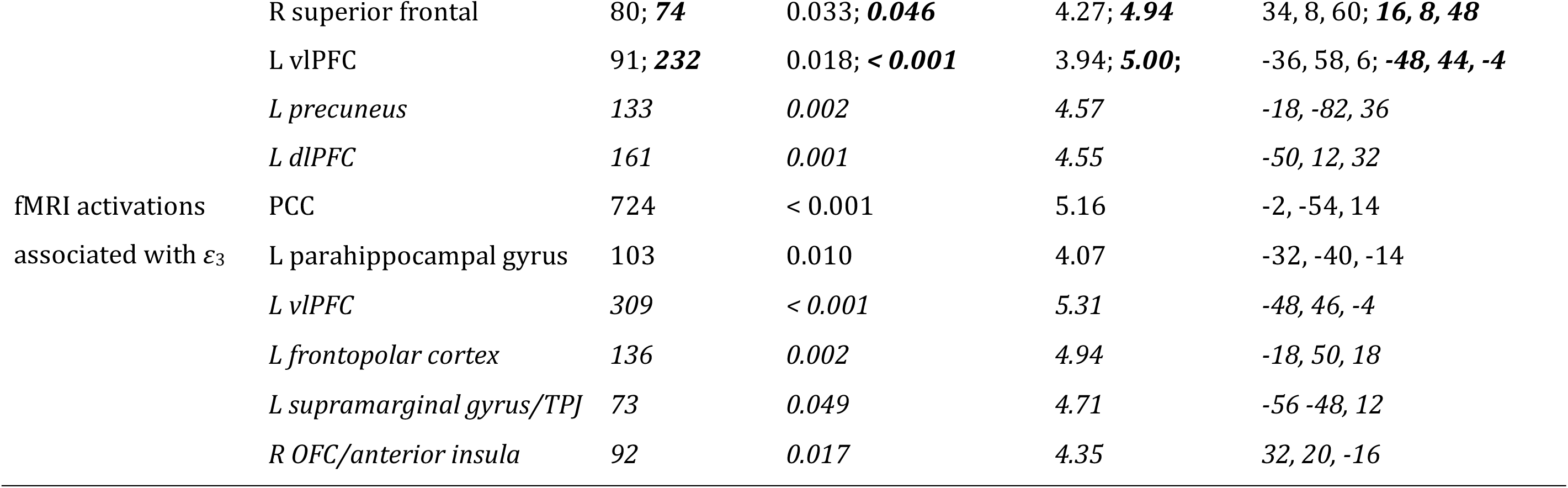
Regions showing main effects (average effect across groups) and group differences of *ε*_2_- (upper panel) and *ε*_3_-related (lower panel) activations during the outcome phase. Upper panel: a contrast-masked (overlapping) subset of the *ε*_2_-related regions display significant group differences (ARMS > controls; **bold** text). Additional regions displaying significant whole-brain corrected group differences are highlighted in *italics* (non-overlapping) or ***bold italics*** (some overlap). Secondary and tertiary values denote these various group difference effects. Lower panel: group differences in *ε*_3_-related activation (controls > ARMS) do not overlap with average effects and thus are shown only in *italics* in distinct rows from the main average effects across groups; dlPFC = dorsolateral prefrontal cortex; OFC = orbitofrontal cortex; PCC = posterior cingulate cortex; TPJ = temporoparietal junction; vlPFC = ventrolateral prefrontal cortex; R = right and L = left hemisphere.

In an additional analysis step, the whole-brain corrected activation pattern was used as a functionally defined mask to restrict the subsequent analysis of group differences in *ε*_2_ (group-by-PE interactions) to regions showing a main effect of *ε*_2_ (note that the two contrasts are orthogonal and thus do not cause non-independence problems for inference). We found significantly greater activation (small volume FWE-corrected p < 0.05) by *ε*_2_ in ARMS patients, relative to controls, in areas of left superior frontal cortex and left anterior insula (Fig. 5B; Table 2). A distinct set of regions also survived whole-brain correction for this contrast, revealing additional group differences in the left vlPFC (some overlap with the pooled whole group region encoding *ε*_2_), precuneus, dlPFC, frontal insulo-opercular cortex, right superior frontal cortex and right anterior insula extending into orbitofrontal cortex (OFC; Fig. 5B; Table 2). An additional region of interest analysis focussing on the same *a priori* midbrain mask described above similarly revealed no significant group differences in activation. Although no significant outcome phase activation was found at the whole-group level for 2^nd^-level uncertainty (*σ*_2_), this variable activated a region of the left fusiform gyrus significantly more in ARMS than in controls (whole-brain cluster-level FWE-corrected; p = 0.030, peak t = 5.44; x = −28, y = −58, z = −10; 82 voxels).

Under whole-brain FWE-correction for multiple comparisons, a number of whole-group average and group difference (controls > ARMS) effects were found for the precision-weighted probability PE (*ε*_3_), which informs updates of beliefs about volatility (Table 2). At the whole-group level, *ε*_3_ significantly activated regions of the bilateral posterior cingulate cortex (PCC) and the left parahippocampal gyrus (Fig. 6A; Table 2). The group contrast ‘*ε*_3_: controls > ARMS’ revealed a more widespread difference in the representation of volatility-related precision-weighted PEs in four additional regions, not overlapping with the whole group average, comprising regions of left vlPFC, frontopolar cortex, supramarginal gyrus/temporoparietal junction and right OFC extending into anterior insula (Fig. 6B; Table 2). Furthermore, within our *a priori* anterior cingulate cortex mask, the subgenual cingulate gyrus (Fig. 6B) displayed significantly greater activation to *ε*_3_ in controls than in ARMS patients (small volume FWE-corrected, p = 0.038, peak t = 4.99; x = −2, y = 16, z = −24; 37 voxels). In addition, an exploratory region of interest analysis using the aforementioned midbrain mask revealed a trend-level cluster (small volume FWE-corrected) of group differences in encoding (*ε*_3_: controls > ARMS) in the left substantia nigra/ventral tegmental area (SN/VTA) of the midbrain (p = 0.075, peak t = 3.54; x = −2, y = −22, z = −20; 5 voxels; Fig. 6B).

**Figure 6.**
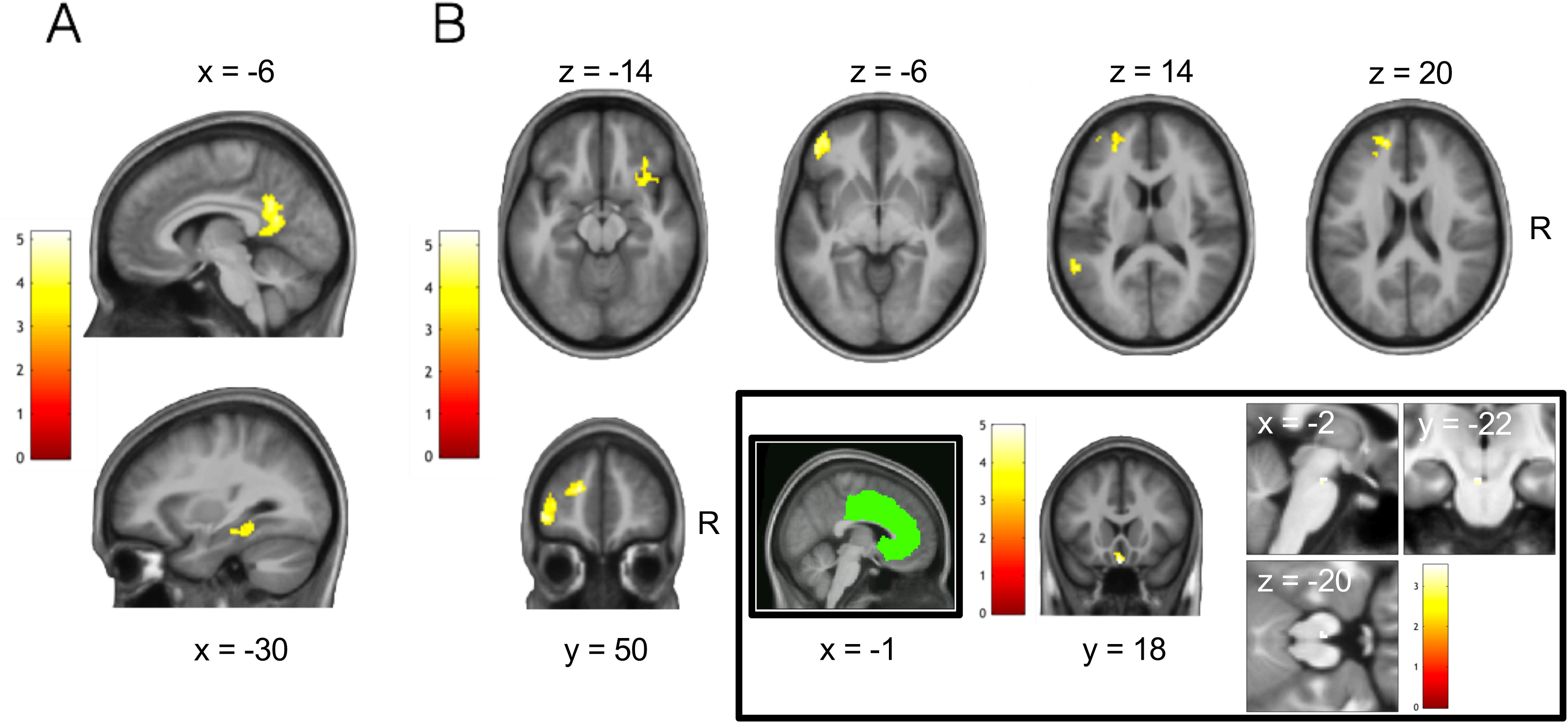
Failures of monitoring and incorporating environmental uncertainty (volatility) in probabilistic learning by ARMS relative to control individuals: prediction error response. (A) A representative map of significant (cluster-level FWE-corrected p < 0.05) group-level (ARMS + controls) outcome-related activations modulated parametrically by precision-weighted volatility-related prediction error (*ε*_3_), calculated via one-sample t-test (N = 25) and overlaid on an anatomical image calculated as the mean structural MRI of the whole group. (B) Greater neural representation of high-level/volatility-related precision-weighted PEs (*ε*_3_) during decision feedback in healthy controls relative to ARMS patients, (main) in a network of regions identified using whole-brain cluster-level correction, (inset) in a region of subgenual anterior cingulate identified under correction for an *a priori* anterior cingulate mask (bottom centre) and in a left midbrain region identified under correction for a dopaminergic midbrain mask (trend level p = 0.075, bottom right). Double inset, green: representative sagittal slice depicting anatomical anterior cingulate cortex mask used as search volume in statistical analysis and multiple comparison correction. Colour bars represent t-statistics.

Finally, when pooling across both groups we found that the predicted volatility 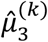 during the decision phase of the probabilistic reversal learning task was associated with BOLD activation in a number of regions including bilateral cuneus, precuneus, superior frontal cortex, right superior temporal and precentral gyrus/central sulcus (Fig. 7A). Contrast masking using this whole group result to examine for group differences revealed two clusters in anterior and posterior superior temporal cortex displaying significantly greater volatility-related activation in controls relative to ARMS (Fig. 7B; Table 3). Two additional regions from a whole-brain corrected group difference contrast, comprising right visual cortex extending into precuneus and a left medial cerebellar region, also showed significantly greater volatility-related modulation of decision-phase activation in controls than in ARMS individuals (Fig. 7B, Table 3). Finally, within our *a priori* midbrain mask we found two regions of SN/VTA, one significant and slightly more dorsal anterior (small volume FWE-corrected p = 0.046, peak t = 3.97; x = 4, y = −22, z = −18; 9 voxels; Fig. 7B) and one trend-level and more ventral posterior (p = 0.075, t = 3.41; −2, −28, −20; 5 voxels; Fig. 7B) that showed greater encoding of predicted volatility in control participants relative to ARMS.

**Figure 7.**
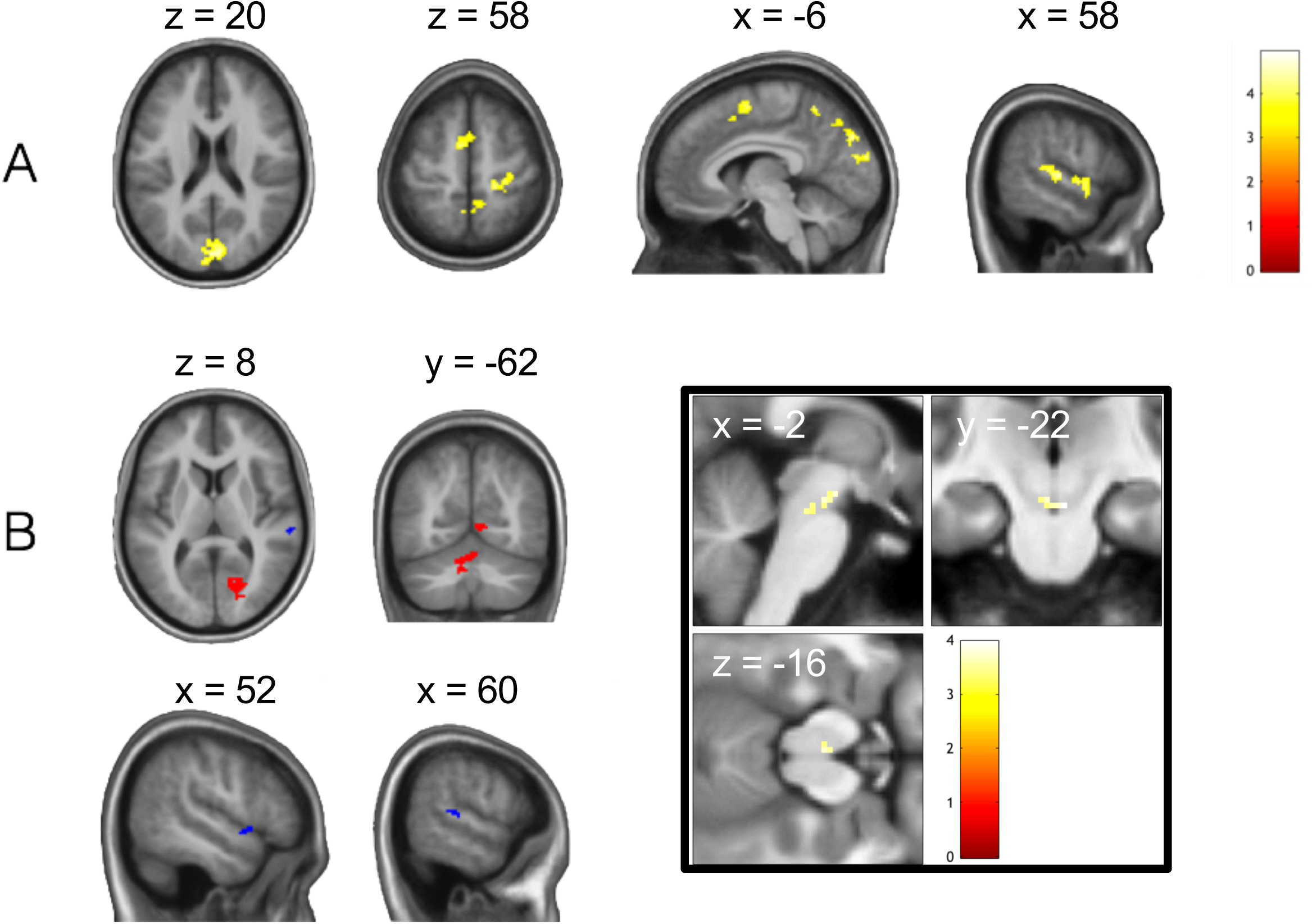
Failures of monitoring and incorporating environmental uncertainty (volatility) in probabilistic learning by ARMS relative to control individuals: decision tracking. (A) A representative map of significant (cluster-level FWE-corrected p < 0.05) group-level (ARMS + controls) decision-related activations modulated parametrically by estimated volatility, calculated via one-sample t-test (N = 25) and overlaid on an anatomical image calculated as the mean structural MRI of the whole group. (B) Greater neural representation of estimated volatility during probabilistic decision-making in healthy controls relative to ARMS patients, (main) in a network where solid colour maps of group differences are binarised and indicate spatial differences between whole-brain cluster-level corrected results (red) and results corrected using the group average map in ‘(A)’ for contrast-masking (dark blue), and (inset) in bilateral midbrain regions identified under correction for a dopaminergic midbrain mask (dorsal cluster p < 0.05, ventral cluster trend level p = 0.051). Colour bars represent t-statistics.

**Table 3.**
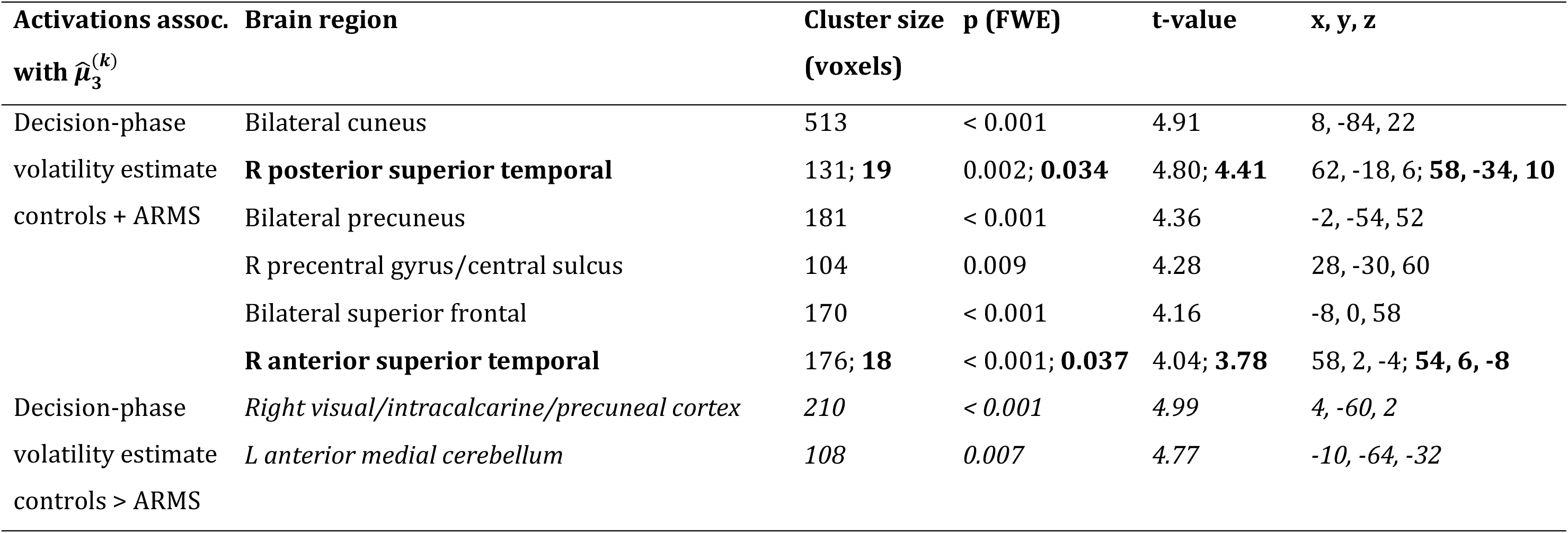
Upper panel: regions showing main effects of estimated volatility-related activation during the decision phase, a subset of which display contrast-masked group differences (**controls > ARMS; bold text**); Lower panel, in *italics*: additional regions showing controls > ARMS differences, identified using whole-brain correction; R = right and L = left hemisphere.

### fMRI association with clinical variables

Following our hypothesis that key clinical features of the ARMS might be associated with the representation of low-level precision-weighted PEs in prefrontal regions, we extracted subject-level fMRI analysis beta-values of *ε*_2_–related activation from the cluster of left superior dlPFC that also displayed a significant group difference in this *ε*_2_ encoding representation (ARMS > controls). We then performed Pearson correlation analysis of these beta-values with sub-scales of the SIPS. We found a significant negative correlation between these fMRI data and the SIPS-GAF measure of current global functioning (r = −0.53, one-tailed p < 0.05; Fig. 5C). In other words, the higher the low-level outcome-related PE encoding in dlPFC, the lower the current functioning and thus the greater the symptom burden or severity at the time of testing.

## Discussion

The ARMS is characterised by attenuated or brief, self-limiting psychotic symptoms, including delusional beliefs. While delusions are thought to reflect the endpoints of aberrant learning and inference processes, with evidence for links to dopaminergic mechanisms in full-blown schizophrenia (Murray et al., 2008; Romaniuk et al., 2010; Gradin et al., 2011), it is not clear whether cognitive disturbances of this sort are already evident in the ARMS.

The specific hypothesis tested in this paper derives from concepts of hierarchical Bayesian inference that play a prominent role in theories of schizophrenia (Stephan et al., 2006, 2009a, Corlett et al., 2009a, 2010; Fletcher and Frith, 2009; Adams et al., 2013). The central idea here is that the brain instantiates a generative model of its sensory inputs, i.e., a model that makes predictions about the environment and how its (hidden) state gives rise to sensations. Perceptual inference rests on inverting the model to determine the most likely cause of sensory input; learning serves to update beliefs such that future sensory inputs can be predicted better. Importantly, under very general assumptions, these belief updates have a generic form: they are proportional to prediction error, weighted by a precision ratio that serves as a dynamic learning rate and balances the expected precision of low-level (e.g., sensory) input against the precision of prior beliefs (see Eqs. 1 and 2; see also Mathys et al., 2014). The corollary from this general update rule is that unusually pronounced belief updates can arise from two sources: by assigning too much precision to sensory inputs, or by overly uncertain prior beliefs.

This perspective allows for formalising the long-standing concept of “aberrant salience” (Kapur, 2003; Heinz and Schlagenhauf, 2010) and predicts that the initial phase of delusion formation could be equally characterised by abnormally high low-level precision (‘sensory precision’) or by abnormally low precision of higher-level, prior beliefs (‘belief precision’). As indicated by Eqs. 1 and 2, these two factors both work in the same direction by elevating the weights of PEs and thus enhancing belief updates. A chronic persistence of either factor could eventually lead to highly unusual beliefs, or necessitate a compensatory adoption of beliefs. For example, constant surprise about events in the world might eventually only be resolved by the adoption of unusual and rigid higher-order beliefs (see, e.g., Corlett et al., 2009b). Conversely, high-order beliefs of abnormally low precision (Adams et al., 2013) render the environment seemingly unpredictable (e.g., more volatile) and boost the weight of lower-level PEs.

Using a combination of computational modelling of behaviour and fMRI, this study examined these hypothetical abnormalities of learning and inference in non-medicated ARMS individuals, compared to control subjects. Firstly, computational modelling of behaviour indicated that, across the duration of the task, ARMS individuals converge to significantly higher levels in their beliefs about volatility. Secondly, concerning the fMRI results, we found evidence for both aberrantly enhanced neural encoding of low-level precision-weighted PEs and aberrantly attenuated encoding of high-level precision-weighted PEs in ARMS individuals.

When testing for the average effect of low-level outcome-related PEs (*ε*_2_) across groups, we found activation in a set of fronto-parieto-insular cortical regions, in line with previous findings (Iglesias et al., 2013; Diaconescu et al., 2017). We found a significant group difference in the degree of this activation within lateral and superior frontal, insular and precuneal cortex, with ARMS individuals displaying increased *ε*_2_-related activation compared with controls. The possibility that this increase in *ε*_2_-related activation might be a prominent pathophysiological characteristic of the ARMS is supported by our observation that beta magnitudes of low-level PE encoding in left superior dlPFC were significantly correlated with more burdensome clinical scores of prodromal symptoms (SIPS-GAF). One mechanistic explanation for the observed overweighting (in terms of neural encoding) of low-level PEs could be a reduced precision of higher-level beliefs in ARMS individuals; in other words, ARMS may be characterised by abnormal estimates of environmental uncertainty.

Accordingly, with respect to activations by higher-level precision-weighted PEs (*ε*_3_) that inform updates of volatility estimates, we indeed found opposing effects, in that many of the same regions – dlPFC, vlPFC and insula, as well as additional effects in temporoparietal junction, the subgenual cingulate and, marginally, the dopaminergic midbrain – displayed greater *ε*_3_–related activations in controls, relative to ARMS individuals. Taken together, our behavioural and fMRI findings related to higher-order beliefs therefore suggest that ARMS individuals may perceive the volatility of the environment differently from controls. The mechanism underlying the observed constellation of an increased ‘set point’ of estimated volatility (according to the behavioural data) and reduced activation in response to high-level precision-weighted PEs (*ε*_3_) that inform updates of volatility estimates, remains to be clarified but suggests a neurobiological difference in the processing of volatility.

Collectively, our results indicate that associative learning under volatility in the ARMS is characterised by higher estimates of environmental volatility (as expressed at the behavioural level) and overly high low-level precision-weighted PE activations (at the neural level). These effects may reflect an enhanced tendency towards belief updating and might explain the empirically observed “jumping to conclusions” bias in ARMS individuals (Broome et al., 2007; Winton-Brown et al., 2015). More generally, this cognitive style may represent a risk factor for delusion proneness.

A final finding is more difficult to interpret: in contrast to the behavioural results, which suggest that ARMS individuals’ updates of volatility were converging to significantly higher levels than those of the controls, fMRI activations by volatility estimates were generally lower in ARMS individuals during decision-making. These relative deactivations by volatility were found in several regions, including temporal and occipital cortices, as well as the midbrain. It is presently not clear how this reduced neural representation of volatility during decision-making is related to the behavioural evidence for increased volatility updating. Generally, however, this atypical cortical representation of volatility does support the general notion that processing of high-level uncertainty is abnormal in ARMS individuals.

Our study has some strengths but also several important limitations that deserve mentioning. Regarding strengths, we implemented strict inclusion criteria that limited recruitment to those ARMS individuals who had never been exposed to antipsychotic medication. Given the potential relation of low-level prediction errors and precision to dopamine function, this step avoided a potentially critical confound. Additionally, we took several steps to maximise the sensitivity of our analyses, including careful correction for physiological noise based on ICA and RETROICOR, orthogonal contrast masking and *a priori* hypotheses about the encoding of specific computational signals in specific anatomical regions of interest. Concerning limitations, while not markedly lower than previous fMRI studies on the ARMS, which have typically examined up to 18 participants (for example, see Allen et al., 2010; Roiser et al., 2013; Schmidt et al., 2013; Falkenberg et al., 2015b; Modinos et al., 2015; Ermakova et al., 2018), our sample size would nonetheless have to be regarded as rather small. We thus emphasise that our results should be treated with caution until replicated in additional samples. A second issue is that the two groups were not perfectly matched with regard to age and sex. We controlled for these differences by including these variables as covariates of no interest in all group-level fMRI analyses. Finally, it is also notable that the SIPS clinical questionnaire we incorporated into our analyses is comprised of a number of different sub-scales, and the reported significant association between one of these (SIPS-GAF) and our fMRI measures of low-level precision-weighted PE encoding would not survive Bonferroni correction for testing all scales. It should thus be treated as a preliminary result that requires confirmation by future studies.

In conclusion, our findings contribute to advancing a mechanistic understanding of cognitive abnormalities during the ARMS. Using computational modelling, functional neuroimaging and clinical measures, we found that behaviour and brain activity of individuals at risk for psychosis, relative to healthy control subjects, shows preliminary evidence for two potential mechanisms – increased low-level precision (of outcome PEs) and greater updates in high-level uncertainty (i.e., volatility) – that converge in their impact and render an individual more prone to adjusting high-order beliefs. In addition, we provide empirical evidence that individual neural representations of outcome-related learning signals (low-level precision-weighted PEs) correlate with individual differences in symptom severity. These findings support previous proposals focusing on the importance of aberrant salience (Kapur, 2003; Roiser et al., 2013) and imprecise higher-order beliefs (Adams et al., 2013) for delusion proneness. The present results may usefully inform future investigations that employ computational and biophysical models to study prediction errors and uncertainty in larger ARMS samples and examine the relevance of these quantities for predicting conversion to psychosis in prospective designs.

## Acknowledgements

We acknowledge support by the University of Zurich, the UZH Clinical Research Priority Program (CRPP) “Molecular Imaging” (DMC, KES), the René and Susanne Braginsky Foundation (KES) and SNF Ambizione PZ00P3_167952 and Krembil Foundation (AOD).

