## Supplementary Materials for "Atypical processing of uncertainty in individuals at risk for psychosis"

#### Methods

*Single-session ICA-based data denoising*

This procedure requires manual labelling of a number of subject datasets in terms of which components resulting from independent component analysis (ICA) are artefactual, in order to train a classifier to automatically perform a denoising analysis on the full dataset. This labelling was conducted by a member of the research team experienced in the identification of artefactual and non-artefactual independent components in functional magnetic resonance imaging (fMRI) data (D.M.C.). Ten participant datasets (5 at-risk mental state, 5 controls) were randomly selected for manual labelling of their ICA results. Based on these labels, the highly accurate FSL FIX machine learning classifier was then trained to distinguish artefact from non-artefact with a conservative (30%) threshold designed to heighten the percentage of components correctly identified as artefactual (although trading off with a theoretical reduction – i.e., one not observed here in practice – in the proportion correctly identified as non-artefactual; see (Salimi-Khorshidi et al., 2014). The trained classifier was then automatically applied to the entire dataset. In the final step of this procedure, denoising was achieved by regressing out the time series of each artefactual component from each subject's preprocessed 4-D fMRI dataset in native space. Prior to statistical analysis, the denoised fMRI data of each subject were transformed into a common space, via linear registration to the individual's high-resolution structural MRI volume using a tissue boundary-based registration (BBR as implemented in FSL 'FLIRT') cost function and then nonlinear warping (as implemented in FSL 'FNIRT') to the MNI152 standard brain template (Montreal Neurological Institute, Canada), with resampling to 2 mm resolution isotropic.

### Table 1a: Prior mean and variance of the perceptual model parameters

**(i) Rescorla-Wagner** $\mathrm{M}_1$

| **Parameter** | **Prior mean** | **Prior variance** |
|---|---|---|
| $\alpha$ | 0.25 | 1 |
| $v^{(k=0)}$ | 0.5 | 1 |

**(ii) Non-hierarchical HGF** $\mathrm{M}_2$

| **Parameter** | **Prior mean** | **Prior variance** |
|---|---|---|
| $\kappa$ | 0 | 0 |
| $\omega$ | -5 | 16 |
| $\vartheta$ | 0.0005 | 0 |
| $\mu_2^{(k=0)}$ | 0 | 0 |
| $\sigma_2^{(k=0)}$ | 1 | 0 |
| $\mu_3^{(k=0)}$ | 1 | 0 |
| $\sigma_3^{(k=0)}$ | 0.1 | 0 |

**(iii) Three-level HGF** $\mathrm{M}_3, \ldots, \mathrm{M}_{10}$

| **Parameter** | **Prior mean** | **Prior variance** |
|---|---|---|
| $\kappa$ | 0.5 | 1 |
| $\omega$ | -5 | 16 |
| $\vartheta$ | 0.5 | 1 |
| $\mu_2^{(k=0)}$ | 0 | 0 |
| $\sigma_2^{(k=0)}$ | 1 | 0 |
| $\mu_3^{(k=0)}$ | 1 | 1 |
| $\sigma_3^{(k=0)}$ | 0.1 | 1 |
| $m$ | 1 | 1 |

Note: The prior variances are given in the space in which parameters are estimated. $\boldsymbol{\kappa}$, $\boldsymbol{\vartheta}$, $\boldsymbol{\alpha}$, $\boldsymbol{\mu_2^{(k=0)}}, \boldsymbol{\mu_3^{(k=0)}}, \boldsymbol{v^{(k=0)}}$ are estimated in logit-space, while $\boldsymbol{\sigma_2}$, and $\boldsymbol{\sigma_3}$ are estimated in log-space.

### Table 1b: Prior mean and variance of the response model parameters

#### Belief to Decision Mapping: Volatility versus Decision Noise

| **Parameter** | **Prior mean** | **Prior variance** |
|---|---|---|
| $\beta$ | 48 | 1 |

Note: The prior variances are given in the space in which parameters are estimated. $\boldsymbol{\beta}$ is estimated in log-space.

Figure S1| **Average Belief Trajectories**

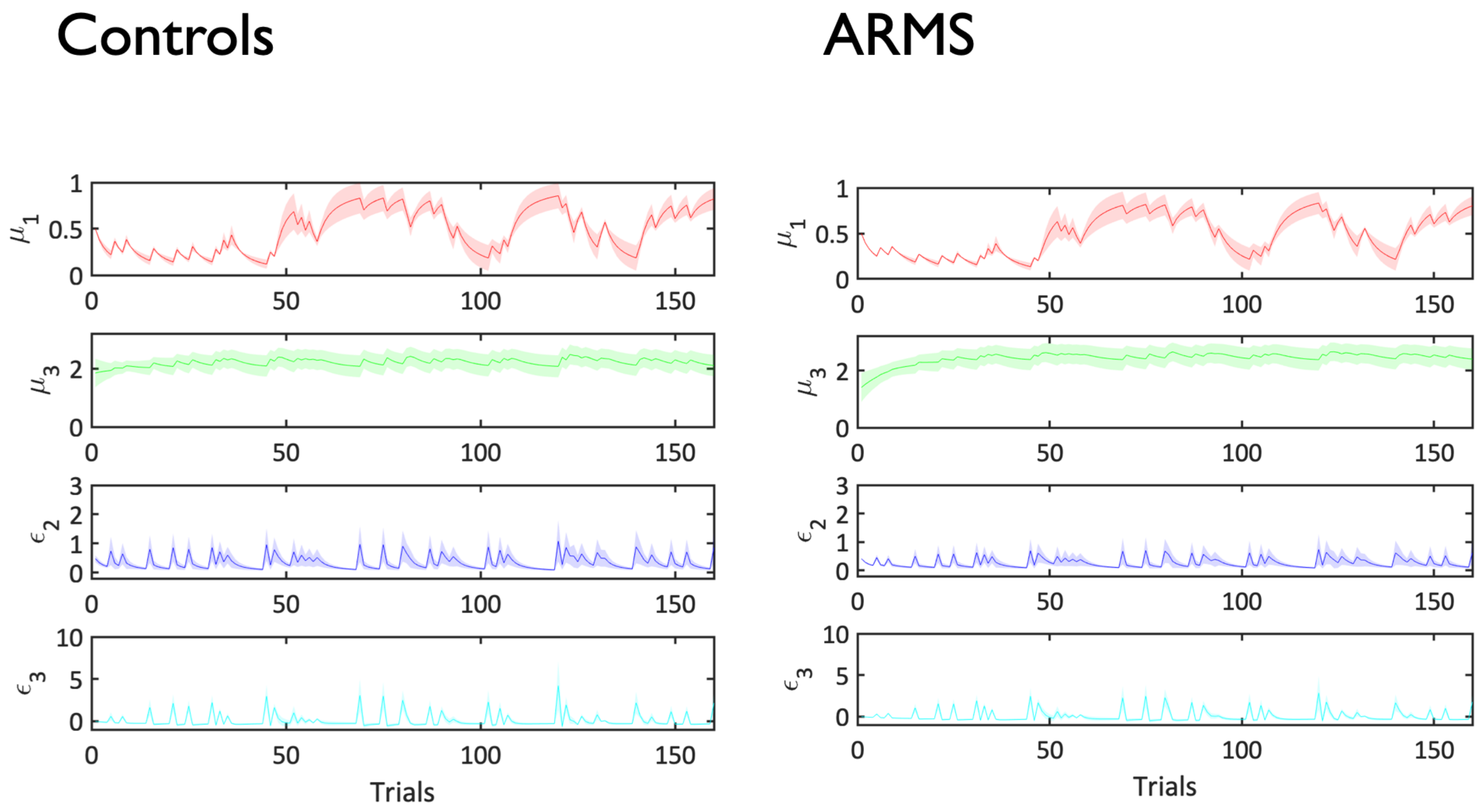


**Figure S1 | Averages of belief trajectories involved in associative learning (predictions and precision-weighted PEs).** This figure includes the average trajectories for the key computational quantities in controls and ARMS. The line plot is generated by averaging the trajectories extracted from the winning model for each participant across trials, i.e., 0 to 160 trials. The shaded area around then depicts +/- standard error of this mean over participants in each group.

### Figure S2| **Individual Belief Trajectories**

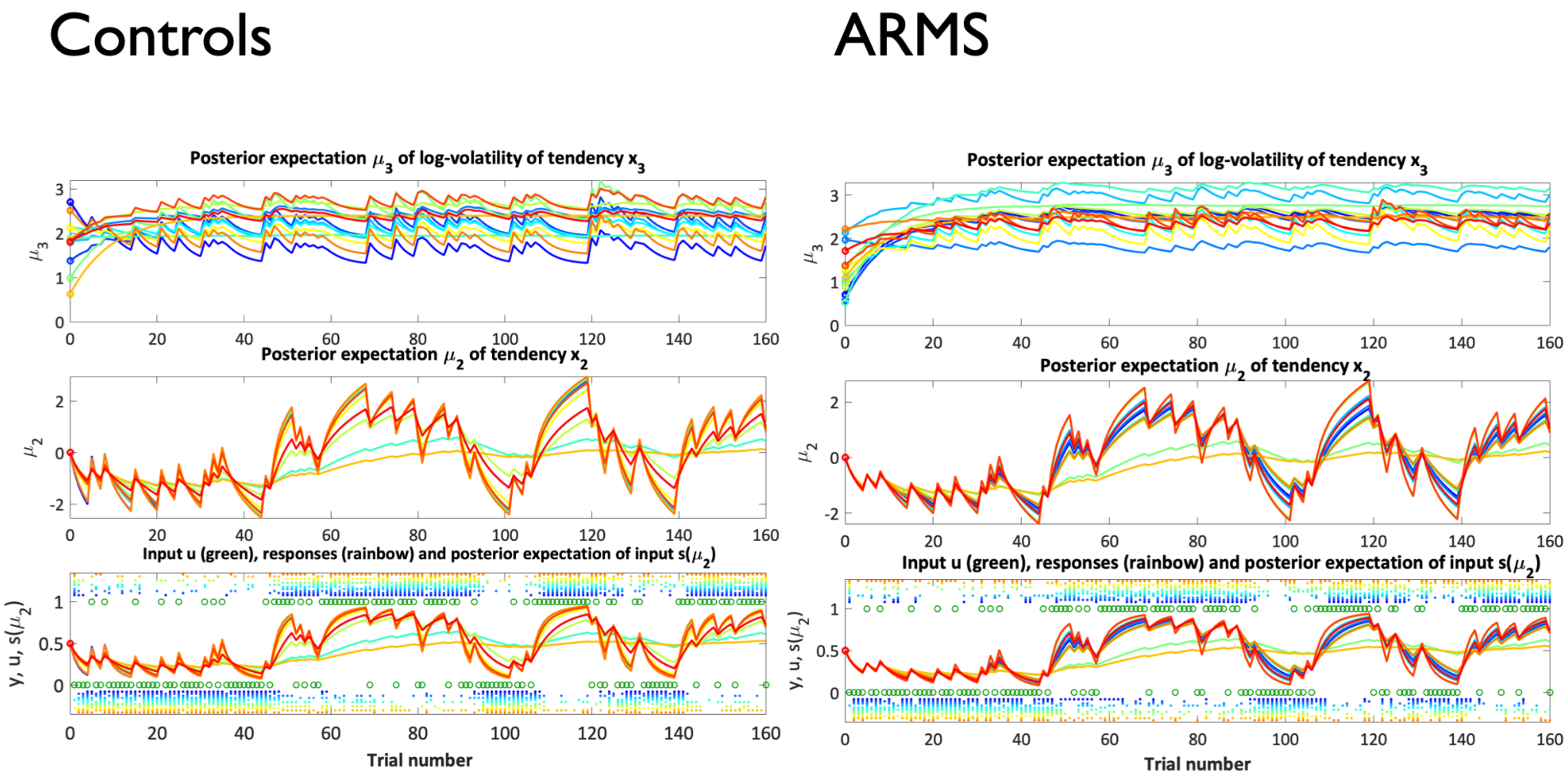


**Figure S2 | Individual belief trajectories.** This figure includes the individual participant belief trajectories at all levels of the hierarchy – beliefs about outcomes, tendency, and volatility – sorted according to subject number for controls and ARMS. At the top, we illustrate each participant's belief about the environmental volatility or $\mu_3$. The middle subplot reflects each participant's estimate of the rewarding cue-outcome association or $\mu_2$, while the bottom graph includes the estimated cue probabilities or $\mu_1$. Here, the green circles above and below the trajectories are the inputs (which were identical across participants). $u = 1$ when a given cue pattern was associated with reward and $u = 0$ when that pattern was unrewarded. The coloured dots above and below the inputs reflect each participant's decision on every trial ($\mathrm{y} = 0$ for a choice consistent with the initially rewarded cue and $\mathrm{y} = 1$ for a choice consistent with the initially unrewarded cue; note that this input structure is 'flipped' relative to Figure 1B in the main manuscript). The dots reflecting the responses are coloured to be identical to the belief trajectories corresponding to each participant.

Figure S3| **Parameter Recovery**

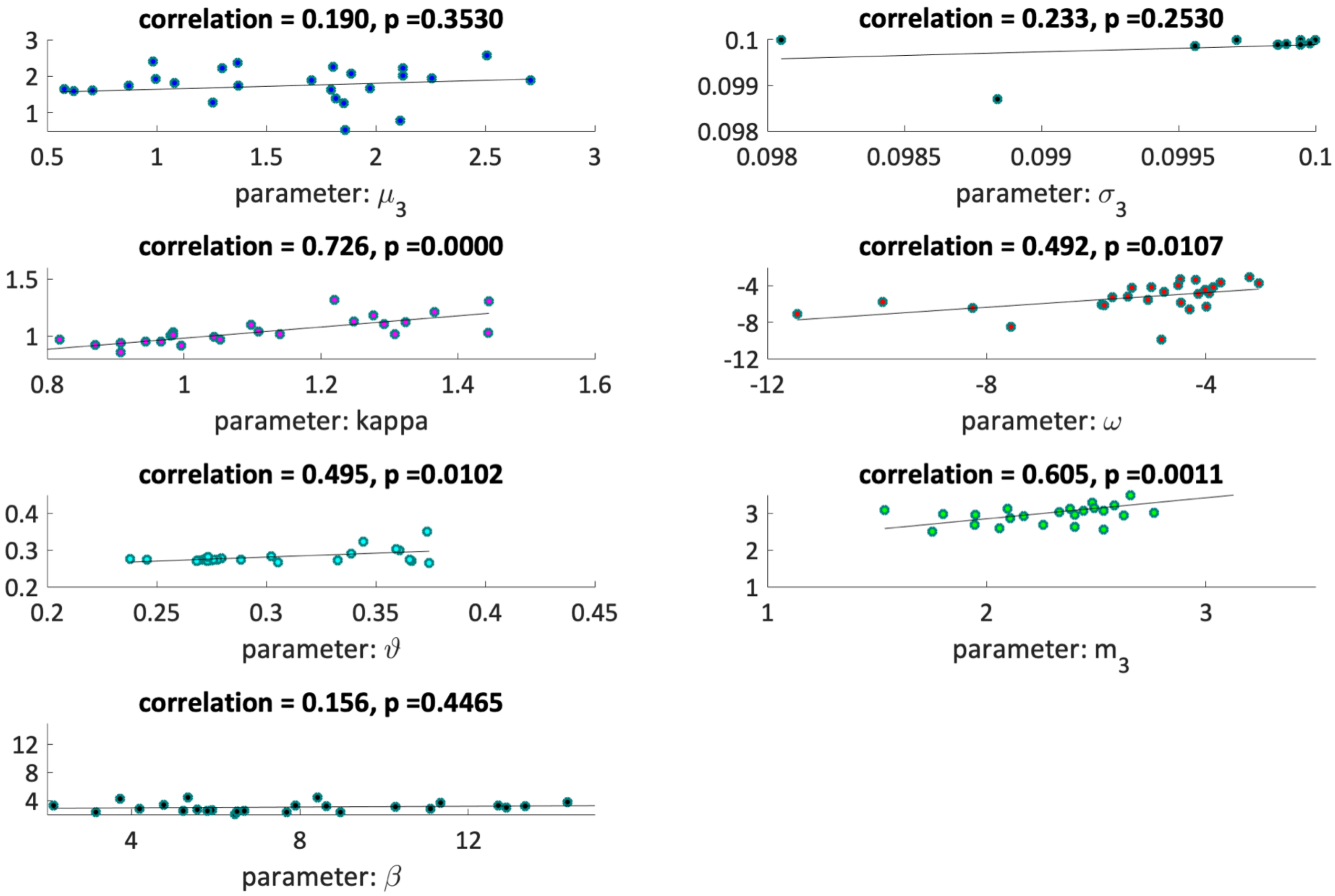

**Figure S3 | Parameter recovery when using empirical parameter values (Mean-reverting HGF).** Parameter recovery for the mean-reverting HGF and parameters $m_3$ (equilibrium value), $\mu_3^{(k=0)}$, $\sigma_3$, $\kappa$, $\omega$, and $\beta$. The correlation coefficients (with corresponding p-values) are included to quantify the parameter recovery results. We saved the seed of the random number generator used in simulations, in order to ensure reproducibility of the results. Please note that, for each plot, we chose to use the same scaling for x- and y-axes.
